# Dopamine transients encode reward prediction errors independent of learning rates

**DOI:** 10.1101/2024.04.18.590090

**Authors:** Andrew Mah, Carla E.M. Golden, Christine M. Constantinople

## Abstract

Biological accounts of reinforcement learning posit that dopamine encodes reward prediction errors (RPEs), which are multiplied by a learning rate to update state or action values. These values are thought to be represented in synaptic weights in the striatum, and updated by dopamine-dependent plasticity, suggesting that dopamine release might reflect the product of the learning rate and RPE. Here, we leveraged the fact that animals learn faster in volatile environments to characterize dopamine encoding of learning rates in the nucleus accumbens core (NAcc). We trained rats on a task with semi-observable states offering different rewards, and rats adjusted how quickly they initiated trials across states using RPEs. Computational modeling and behavioral analyses showed that learning rates were higher following state transitions, and scaled with trial-by-trial changes in beliefs about hidden states, approximating normative Bayesian strategies. Notably, dopamine release in the NAcc encoded RPEs independent of learning rates, suggesting that dopamine-independent mechanisms instantiate dynamic learning rates.

## Introduction

Reinforcement learning describes how animals or agents learn the value of states and actions to select actions that maximize future expected rewards^1^. Reinforcement learning algorithms, including temporal-difference learning, update state and action values using reward prediction errors (RPEs), or the difference between experienced and expected rewards. The rate of errordriven learning is often assumed to be constant, but work across humans, monkeys, rats, and mice, has found behavioral evidence for dynamic learning rates^2–8^. In volatile environments, dynamic learning rates allow animals to learn faster when the world is changing, and slower when the world is stable^9–11^.

Dopaminergic inputs to the striatum, the input structure of the basal ganglia, are thought to convey a biological RPE. In reinforcement learning models of the basal ganglia, cortical inputs to the striatum convey the animal’s state, with the strength of the synapse proportional to the expected future reward, or value, of that state^1,12,13^. States with higher values have stronger synapses that are more likely to drive striatal action selection. The strengths of these corticostriatal synapses are updated via dopamine-dependent plasticity, proportional to dopamine RPEs^14–17^. However, reinforcement learning algorithms update values proportional to the *product* of the RPE and learning rate. It is currently unclear whether dopamine conveys only the RPE, with other substrates dictating the learning rate, or whether dopamine encodes the RPE scaled by the learning rate. These scenarios are indistinguishable if learning rates are static. Here, we leveraged the fact that reinforcement learning is dynamic in changing environments to characterize dopamine encoding of learning rates in the nucleus accumbens core (NAcc) by recording dopamine release in rats performing a task with latent reward states.

## Results

### Rats use a dynamic learning rate

We trained rats on a self-paced temporal wagering task with semi-observable reward blocks^18^(Fig. 1A-B). Rats were offered different volumes of water rewards (5, 10, 20, 40, 80*µ*L), cued by an auditory tone. On 75-85% of trials, rewards were delivered after variable, unpredictable delays drawn from an exponential distribution. On 15-25% of trials, rewards were withheld. The rats could choose to wait for the water reward, or could opt-out at any time to start a new trial. We introduced uncued blocks of trials with differing reward statistics: low blocks, which offered the smallest three rewards (5, 10, 20*µ*L), and high blocks, which offered the largest three rewards (20, 40, 80*µ*L), were interleaved with mixed blocks, which offered all rewards (Fig 1B).

**Figure 1:**
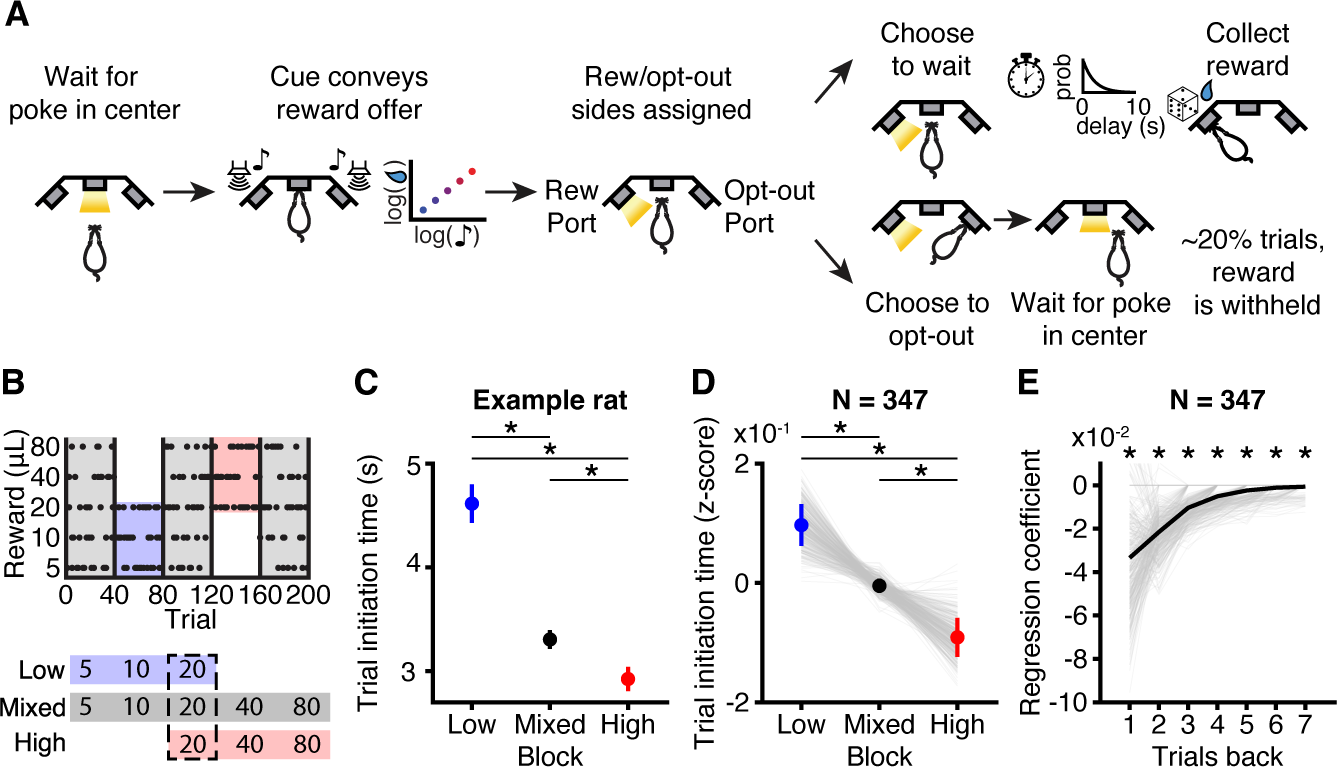
Trial initiation times are sensitive to previous rewards and blocks. **A.** Task schematic. **B.** Block structure. **C-D.** (C) Average trial initiation times for a single rat (p*<*0.01, Wilcoxon rank-sum test) and (D) all rats (*p<<*0.001, Wilcoxon signed-rank test, *N*=347) are sensitive to block. **E.** Regression coefficients of previous rewards predicting trial initiation times across rats (Asterisk indicate *p<*0.05, Wilcoxon signed-rank test, *N*=347). All error bars are mean*±*S.E.M.

We measured the time between the rat’s final poke in the reward or opt-out port and the start of the next trial (trial initiation time). Trial initiation times were inversely proportional to the value of the environment, and provided a continuous behavioral readout of rats’ estimates of state values^18^. Rats were slower to initiate trials in low blocks compared to high blocks (Fig. 1C-D). Furthermore, when we regressed initiation times against previous reward offers, coefficients were larger for more recent offers, and gradually decreased for more distant trials (Fig. 1E). This pattern is consistent with canonical reinforcement learning algorithms which estimate the value of the environment as a recency-weighted average of previous rewards. We focused on trial initiation times because our previous work found that initiation times reflected state value estimates that were updated consistent with standard reinforcement learning algorithms, whereas willingness to wait during the reward delay reflected more complex state inference processes that are distinct from these standard algorithms^18^.

In reinforcement learning, the decay time of previous trial coefficients is directly proportional to the learning rate parameter, which is often assumed to be static. However, examination of trial initiation times aligned to transitions from low or high blocks into mixed blocks revealed two phases of learning- an initial phase of fast learning, followed by slower dynamics later in the block, suggestive of higher learning rates immediately following block transitions (Fig. 2A; Supp. Fig. 1). Previous work found that the overshoot in trial initiation times after block transitions could not be explained by a static learning rate^18^, and is robust across rats (Supp. Fig. 2). Consistent with this result, when we regressed trial initiation times against previous rewards separately for the first and last ten trials of each mixed block, rats integrated over fewer trials earlier in the block, i.e., had higher learning rates, compared to later in the block (Fig. 2B-C; Supp. Fig. 3-4). Across rats, exponential functions fit to the regression coefficients had significantly smaller time-constants (higher learning rates) for early versus late trials (Fig. 2D).

**Figure 2:**
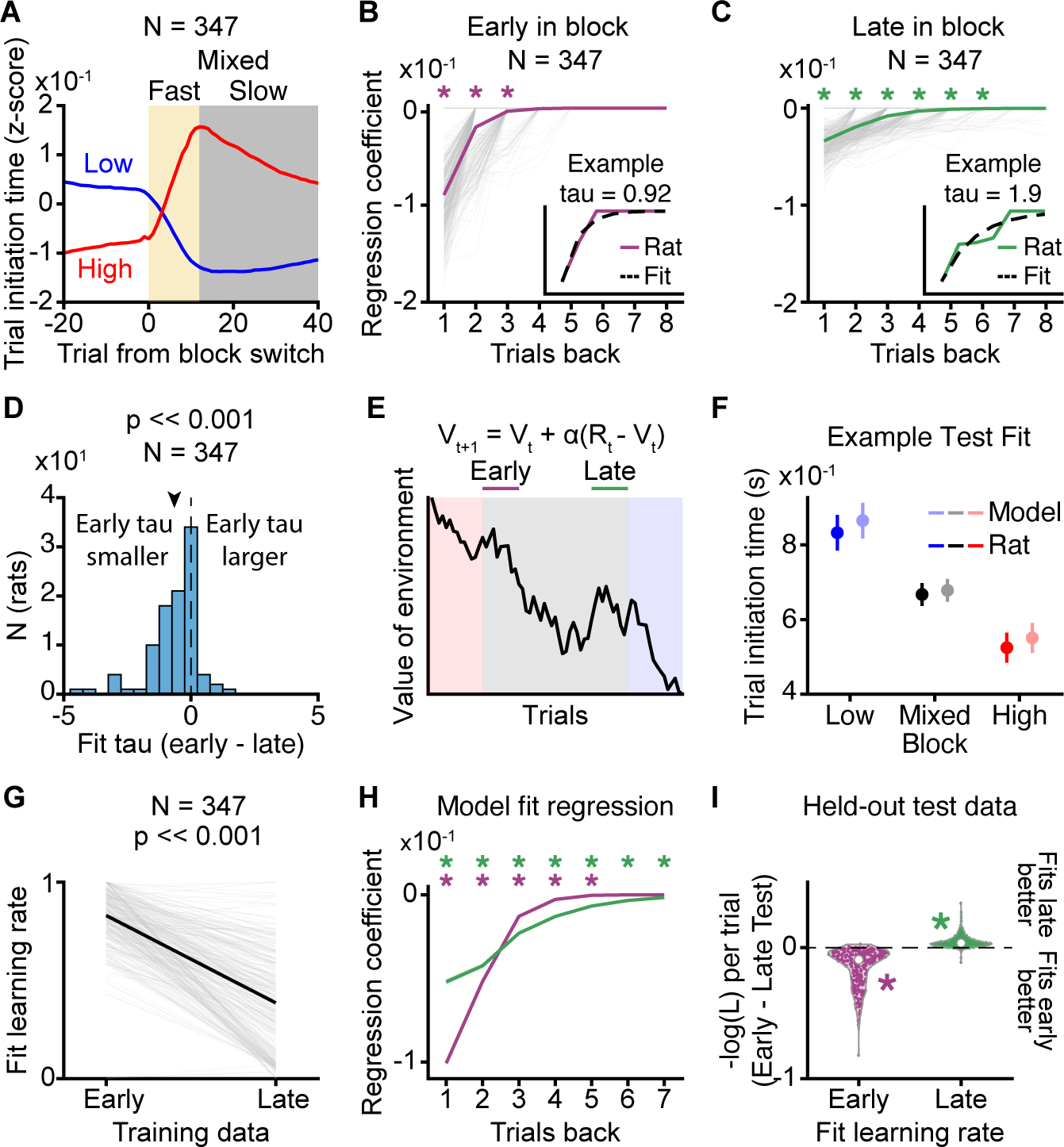
Rats use a higher learning rate early in mixed blocks. **A.** Trial initiation times aligned to transitions into mixed blocks from low (blue) and high (red) blocks. Smoothed with a causal filter of size 10 trials. **B-C.** Previous reward regression coefficients for (B) early and (C) late mixed block trials (Wilcoxon sign-rank test, *N*=347). Insets show example regression coefficient (solid) overlaid with exponential fit (dashed) **D.** Difference in time-constant, tau, of exponential decay fit to early and late mixed block regression coefficients across rats. Arrow indicates mean (*p<* 0.001, paired Wilcoxon Sign-rank test, *N*=347). **E.** Model schematic. **F.** Example model performance on held-out late trial test data. **G.** Recovered learning rate parameters for early and late mixed block training data across rats (*p<* 0.001, Wilcoxon Sign-rank test, *N*=347). **H.** Average previous trial regression from model fits to early (purple) or late (green) trials across rats (*N*=347). **I.** Comparing negative log-likelihood of early and late parameters on held-out test data. White circles indicate mean. (Wilcoxon sign-rank test, *N*=347). Asterisks indicate *p<* 0.05. Arrow bars are mean*±*S.E.M.

We next fit a simple reinforcement learning model with a static learning rate separately to early and late trials (Fig. 2E). The model estimated the value of the environment according to the recursive equation: *V_t_*_+1_ = *V_t_* + *α*(*R_t_ − V_t_*), where *R_t_ − V_t_*is the RPE, and *α* is a learning rate that dictates the dynamics with which values are updated over trials. The trial initiation times were modeled as inversely proportional to this value, which captured the rat’s behavior on held-out test data^18^ (Fig. 2F). Models fit to early mixed block trials had significantly higher learning rates (Fig. 2G) and fewer significant previous trial coefficients (Fig. 2H) compared to models fit to late mixed block trials. Model comparison confirmed that learning rates fit to early and late parts of the block provided better fits to held out test data in early and late block trials, respectively (Fig. 2H). Overall, these data suggest that rats use a dynamic learning rate that is higher following block transitions compared to later in the block.

### Rats’ dynamic learning rates reflect changing beliefs about reward blocks

Previous work found that animals adjust their learning rates depending on the volatility in the environment, since it is advantageous to learn faster in dynamic environments^3,5,7^. To determine which features of the environment the rats were using to adjust their learning rates, we tested several classes of dynamic learning rate models (Fig. 3A). Previous studies have proposed that subjects might scale their learning rates based on salient events or outcomes^19,20^. We modeled salience as proportional to the log-reward offer on each trial, so larger rewards were assumed to be more salient than smaller rewards, which we call the Mackintosh model, following [19].

**Figure 3:**
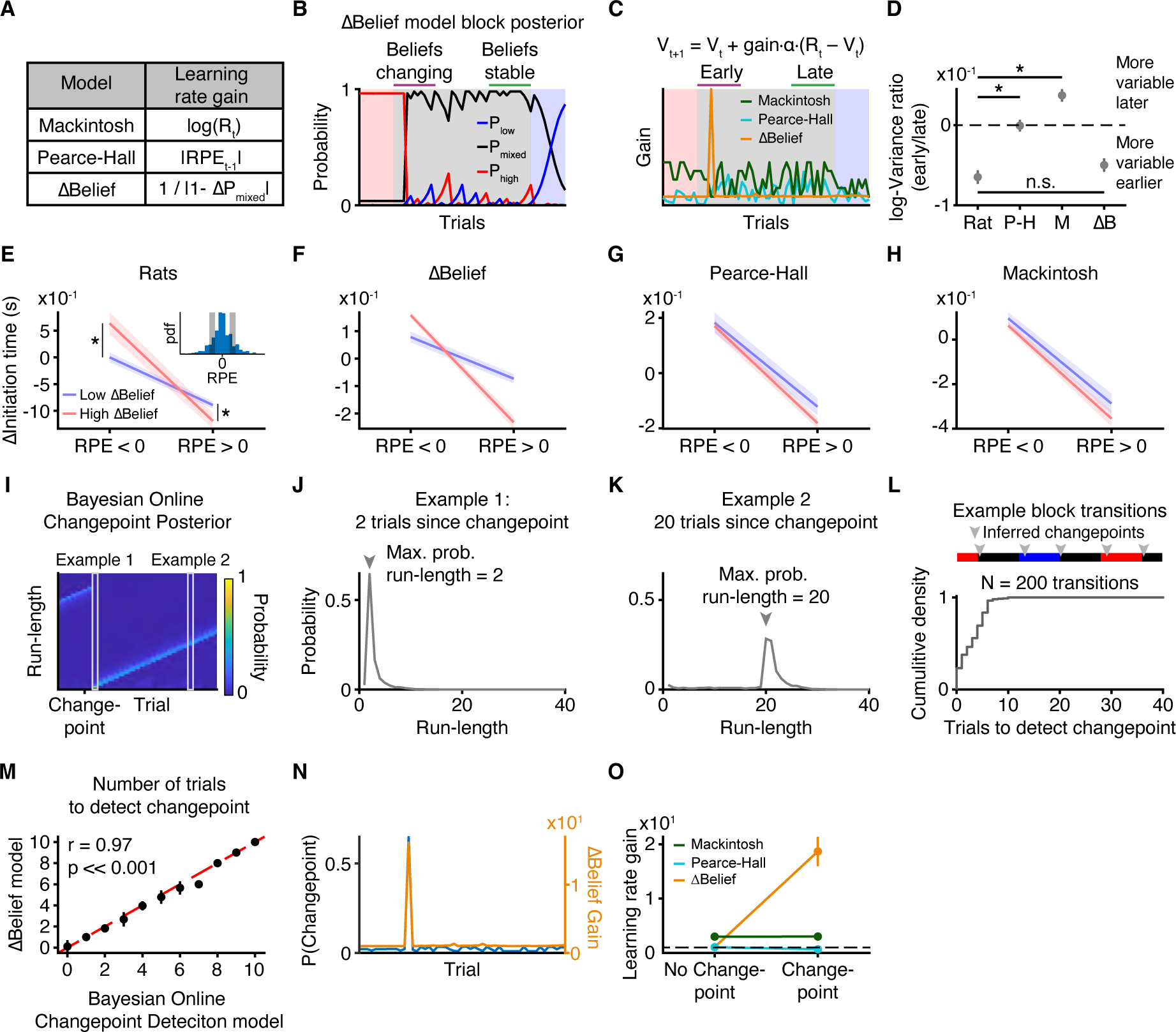
Rats use changing beliefs to modulate learning rates. **A.** Explanation of models. **B.** Block posterior probability from ΔBelief model. Beliefs change the most at block transitions. **C.** Examples of learning rate gain from (A). **D**. Trial initiation time variance early vs. late for dynamic learning rate models and rat data. (Wilcoxon rank-sum test, *N*=347). **E.** Change in trial initiation time for negative and positive RPE bins (inset) for large (red) or small (blue) change in mixed block belief (*>* or *<*50th percentile; Wilcoxon sign-rank test, *N*=347). **F-H**. Model predictions for (E) ΔBelief, (F) Pearce-Hall, and (G) Mackintosh model. **I.** Bayesian online changepoint detection model run-length posterior over trials around block transitions. **J-K.** Example changepoint posteriors for a trial (J) two trials from a changepoint or (K) 20 trials from a changepoint. Arrows indicate trial with maximum posterior probability. **L**. Simulated changepoint posterior for Bayesian Online Changepoint Detection model (Top) Colored rectangles indicate true block, and triangles show inferred changepoints. (Bottom) Cumulative probability distribution for number of trials between block transitions and inferred changepoint (*N*=200 simulated transitions). **M.** Average number of trials to detect changepoint from ΔBelief model and Bayesian Online Changepoint Detection model (*N*=200 simulated transitions). **N.** ΔBelief model learning rate gain correlates with changepoint probability from Bayesian Online Changepoint Detection model. **O**. Learning rate gain for dynamic learning rate models on inferred changepoint trials. Asterisks indicate *p<* 0.05. Error bars are mean*±*S.E.M.

Alternatively, previous work suggests that it is advantageous to modulate learning rates based on perceived volatility in the environment^3,5,21^. When volatility is high, previous outcomes are less predictive of future states, so agents should use a higher learning rate to disregard distal outcomes. We tested two classes of volatility-based dynamic learning rate models. First, the model-free Pearce-Hall model adjusts the learning rate proportional to the unsigned reward prediction error of the previous trial^22^. Intuitively, large RPEs indicate that reward expectations are inaccurate, potentially because the environment is changing. Second, we characterized a “model-sensitive” model we previously developed^18^, which we refer to as the ΔBelief model. This model uses Bayes’ rule to infer the probability of being in each block given the rat’s current and past reward offers, and scales the learning rate by the trial-by-trial change in the belief about the block (i.e., change in posterior probability; Methods).

We used these above models to generate qualitative predictions about initiation time behavior. First, the models make distinct predictions about how the variance of the initiation times should change within mixed blocks. Higher learning rates imply higher variability in initiation times because the value estimate is updated more on each trial. Because the mixed blocks include all reward offers, the magnitude of RPEs is comparable both early and late in the blocks. Therefore, the Pearce-Hall model, which scales the learning rate by the previous unsigned RPE, predicts similar learning rates early and late in mixed blocks, and thus similar initiation time variance (Fig. 3C). Similarly, large rewards are equally likely early and late in mixed blocks, so the Mackintosh model, which scales the learning rate with the log-reward offer, predicts similar initiation time variance across mixed blocks (Fig. 3C). However, the ΔBelief model, which scales the learning rate with the trial-by-trial change in the belief about the inferred block, predicts higher variance in initiation times early in the block when beliefs are changing, compared to late when beliefs are stable (Fig. 3B-C). We measured the variance of the initiation times for the first and last ten trials of each mixed block, and computed their log ratio (negative values correspond to higher variance early in the block). Consistent with the ΔBelief model, but not the other models, the variance of rats’ initiation times was higher in the early mixed block trials, compared to later trials (Fig. 3D).

Next, we explicitly tested a prediction from the ΔBelief model - that rats’ trial-by-trial learning should depend on the change in belief about the block, controlling for RPEs. We focused on early trials in the block, when there should be a broader range of beliefs to increase statistical power. We estimated RPEs using model parameters fit to behavior from the first ten trials of each block. We compared the trial-by-trial change in initiation times for the same values of RPEs, but conditioned on whether the ΔBelief on that trial was high or low (*>* or *<*50th percentile). We found that rats changed their initiation times more for both positive and negative RPEs on trials with large changes in beliefs, compared to trials with the same binned RPEs but small changes in beliefs (Fig. 3E), consistent with the ΔBelief model (Fig. 3F), but not the other dynamic learning rate models (Fig. 3G-H). In summary, rats use knowledge about the underlying task structure to update their learning rates.

However, it remains unclear why rats would use the ΔBelief model. We found that the ΔBelief model approximates a normative algorithm known as Bayesian online changepoint detection^23^. Bayesian online changepoint detection uses sequential observations (e.g., reward offers) and a model of the environment to estimate the probability that the underlying distribution generating those observations (e.g., the reward block) has changed. These inferred transitions are called changepoints. Specifically, on each trial, the model generates a distribution over the number of trials since the last changepoint (run-lengths) and chooses the run-length that maximizes the posterior probability (Fig. 3I-K). A run-length of N means the model estimates that a changepoint occurred N trials ago (Fig. 3J-K). While Bayesian online changepoint detection can correctly estimate changepoints around block transitions in our task (Fig. 3L), this model is computationally expensive. On each trial, the model iterates over all previous trials to find changepoints- the number of computations is a quadratic function of number of trials (Methods). Strikingly, we found that the gain term from the ΔBelief model, which is less computationally expensive, is highly correlated with the changepoint probability (Fig. 3M). Over many simulated sessions, the ΔBelief gain was significantly higher on trials with inferred change-points compared to other trials. There was no systematic relationship between the unsigned RPE (Pearce-Hall model), or log reward offer (Mackintosh model), and inferred changepoints (Fig. 3N-O). The ΔBelief model can therefore provide a simple approximation of a normative changepoint detection model that allows rats to detect changes in the environment and adjust their learning accordingly, similar to previous work that found behavioral evidence for simpler approximations of changepoint detection^24^.

### Dopamine release in the NAcc is not modulated by the dynamic learning rate

We next sought to find neural correlates of the dynamic learning rate in the NAcc, where dopamine is thought to mediate trial-by-trial learning by instantiating a biological RPE^25–38^. Mechanistically, dopaminergic RPEs are thought to mediate plasticity at synapses onto medium spiny neurons, the principal cells of the striatum, to increase (or decrease) the likelihood of taking actions in a state that previously produced positive (or negative) RPEs. This account of dopamine function implicitly assumes dopamine release represents the product of the learning rate and the RPE, and so should reflect dynamic learning rates in our task.

We focused our recordings on the NAcc, which is thought to determine the vigor of motivated behaviors^39^. Recent work from our lab has also found that dopamine RPEs in the NAcc causally determine initiation times on subsequent trials^40^. We recorded dopamine release in the NAcc (*N*=14, Supp. Fig. 4) using fiber photometry and GRAB_DA_, a fluorescent dopamine sensor. We observed robust phasic dopamine responses at the time of the offer cue that were consistent with an RPE. First, an RPE should correlate with reward offer. We found that NAcc dopamine release scaled monotonically with offered reward volume in mixed blocks, with dips for smaller rewards (Fig. 4B,D). An RPE signal should also scale inversely with expectations. Focusing on 20*µ*L trials, which appeared in all blocks, dopamine increased on 20*µ*L trials in low blocks, when reward expectations were low, and decreased on 20*µ*L trials in high blocks, when expectations were high (Fig. 4C,E). Third, when we regressed NAcc dopamine release (area under the curve from 0 to 0.5 seconds) against reward history, we found positive coefficients for the current trial and negative coefficients for previous trials, a hallmark of RPE encoding^29^ (Fig. 4F). Finally, NAcc dopamine release correlated with RPEs estimated from the behavioral model fit separately to the first and last 10 trials of each block (Fig. 4G,I).

**Figure 4:**
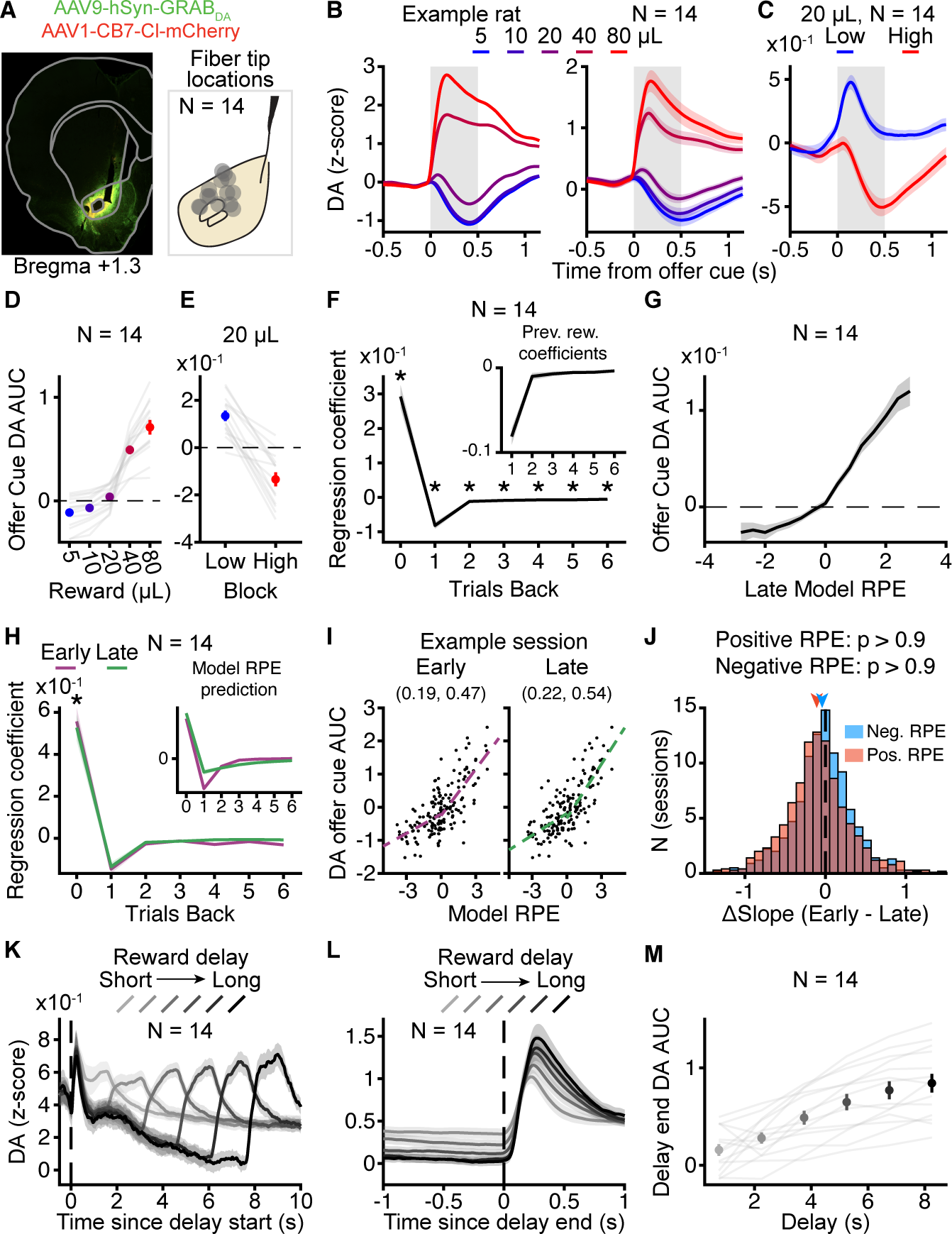
Dopamine release in the NAcc encodes reward prediction errors independently of learning rates. **A.** Example NAcc histology and recording site summary. **B.** NAcc dopamine aligned to offer cue by rewards in mixed blocks for an example rat (left) and averaged across rats (right; *N*=14). **C.** Average NAcc dopamine response to offer cue for 20*µ*L by block (*N*=14). **D.** Average AUC from B. (gray box, *N*=14). **E.** Average AUC from C. (gray box, *N*=14). **F.** Reward history regression coefficients for NAcc dopamine signal. Inset shows previous reward coefficients (Sign-rank test, *N*=14). **G.** Offer cue AUC by binned late model RPE (*N*=14). **H.** Reward history regression coefficients for NAcc dopamine in early (purple) and late (green) trials. Paired sign-rank test, *N*=14. Inset shows model prediction regression RPE vs. previous rewards for parameters fit to early vs. late trials. **I.** Example rat NAcc dopamine response by binned RPE for early (left) and late (right) trials (solid) with positive and negative RPE regressions (dashed). Parentheticals indicate positive and negative slopes, respectively. **J.** Difference between early and late regression slopes for DA for positive (orange) and negative RPEs (blue) over individual sessions for all rats. Arrows indicate mean (*N*=994 sessions, *p*_early_,*p*_late_ *>* 0.9, one-tailed Wilcoxon sign-rank test. **K.** NAcc dopamine aligned to beginning of delay period for rewarded mixed block trials by reward delay, *N*=14. **L.** NAcc dopamine aligned to end of delay period for rewarded mixed block trials as a function of reward delay, *N*=14. **M.** Baseline-corrected average AUC from L., *N*=14. Asterisks indicate *p<* 0.05. Error bars are mean*±*S.E.M.

Next, we compared NAcc dopamine release early vs. late in the block when the rats used different learning rates. If dopamine release is the product of the RPE and the learning rate, then dopamine should be influenced by fewer trials in the past when the learning rate is higher (Fig. 4H insert). However, there was no significant difference between previous trial regression coefficients fit to NAcc dopamine release early or late in the block (Fig. 4H). This hypothesis also predicts that dopamine release would be greater for the same RPE earlier in the block, when the learning rate is higher, compared to later in the block, when beliefs are stable and learning rates are lower. To account for nonlinear encoding of RPEs, we fit separate regression lines to dopamine encoding (AUC) of positive and negative model RPEs for each session. There was no significant difference in the slope parameters fit to early or late trials in each session across rats, and this was true for both positive and negative RPEs (Fig. 4I,J; Supp. Fig. 5). Therefore, despite strong behavioral evidence for higher learning rates earlier in mixed blocks when beliefs about hidden states are changing, we did not find differential NAcc dopamine dynamics between early and late block trials.

Previous work found that the activity of VTA dopamine neurons can reflect beliefs about hidden task states if rewards are probabilistic and variable in their timing^41^. We therefore examined NAcc dopamine during the delay period. Because rewards were omitted on a subset of trials, as animals waited for the reward, there is ambiguity about whether they are in a rewarded or unrewarded trial. Sorting rewarded trials based on delay duration revealed negative dopamine ramps during the delay (Fig. 4K), consistent with previous studies^41^. These negative ramps were also apparent as different baselines when the data were aligned to the cue indicating that reward was available (Fig. 4L). These ramps have been interpreted as moment-by-moment negative RPEs as the animals wait without receiving a reward^32^. When a cue indicated that reward was available, there was a large phasic dopamine response in the NAcc, the magnitude of which scaled with the delay (Fig. 4K-M). This pattern cannot be captured by traditional model-free temporal difference learning, and requires additional knowledge about the task structure^41^- the probability of being in an unrewarded trial increases over time, so the reward cue is more unexpected given beliefs about the trial. Therefore, NAcc dopamine reflects beliefs about hidden states at the time of probabilistic rewards, but not about latent reward blocks at the time of the offer cue.

## Discussion

Our results offer a nuanced, “model-sensitive” view of dopamine activity, in which different aspects of state inference differentially modulate dopamine release in the NAcc. On one hand, NAcc dopamine at offer cues encodes RPEs independently of inference about the latent reward block, despite beliefs about the blocks strongly influencing reinforcement learning and behavior (Fig. 4H-J). This is consistent with traditional model-free interpretations of dopamine activity. However, at the end of the delay period, we found delay-dependent patterns of NAcc dopamine release that cannot be explained by model-free temporal difference learning, consistent with previous work^41^ (Fig. 4K-M). These results add to a body of work expanding dopamine beyond traditional, model-free learning algorithms^35,42^.

It remains unclear why some aspects of state inference modulate NAcc dopamine while others do not. In a conditioning paradigm with probabilistic rewards, during the delay period, there is ambiguity about the state the animal is in: whether it is in the delay period, or if it has transitioned to the inter-trial interval on an unrewarded trial. In other words, there is *uncertainty* about the current state. However, this type of uncertainty is qualitatively different from the uncertainty about the latent reward blocks in our task. Uncertainty in the timing of probabilistic rewards reflects *expected*, or irreducible, uncertainty^21^. Even with a perfect model of the world, there is inherent uncertainty in the outcome of stochastic events - one can know the probabilities of a coin toss, but cannot predict the outcome of the next flip. This contrasts with *unexpected* uncertainty, which is related to the volatility of the environment - a perfect model of the world now may not be perfect if the world changes^21,43^. In other words, unexpected uncertainty about the reward blocks can be reduced with additional observations, while expected uncertainty about probabilistic reward timing is irreducible. We find that dopamine release in the NAcc reflects expected but not unexpected uncertainty. Together, these results suggest that different types of uncertainty may map onto neurobiologically distinct mechanisms, with potentially dissociable consequences for learning^8,21,34,43–45^.

Previous work has suggested that dopamine release in the NAcc directly encodes the learning rate^46^, implicating dopamine in a broader class of policy learning algorithms, beyond traditional value learning. However, our work examines distinct phases of learning from [46]. We exclusively recorded from expert, as opposed to naive, animals. Presumably our expert rats have already learned their final behavioral policy, and must only learn about the current state of the environment. By contrast, task naive animals are simultaneously learning the associative structure of the environment and how to optimally behave in that environment. Such distinct learning goals likely engage NAcc dopamine differently and could explain the differences in our findings.

One key future question is what could be driving the dynamic learning rate at the level of synaptic plasticity. Previous work has shown that plasticity at corticostriatial synapses depends on the coordinated activity of dopamine^14–17^ with other neuromodulators, like acetylcholine^47^ and serotonin^48–50^. Serotonin neurons, which project to the NAcc from the dorsal raphe nucleus^51^, have been shown to encode unexpected uncertainty^7^, and can causally influence learning rates in mice^52^. Other neuromodulators that have been hypothesized to encode unexpected uncertainty, like norepinephrine^43^, do not strongly project to the NAcc^53–55^, but could influence learning in other neural circuits. Future studies clarifying task-related dynamics of other neuromodulators may elucidate the circuit mechanisms that combine dopaminergic RPEs with trial-by-trial changes in beliefs to modulate the rate of learning at behavioral and synaptic levels.

### Limitations of the study

Our current study focused on recording dopamine release in the NAcc. However, given recent findings that show considerable heterogeneity in dopamine activity across the striatum^56–67^, additional studies are required to understand how dopamine activity across striatal subregions is modulated by hidden-state inference. More specific recording techniques, such as optogenetically-tagged recordings or cell-type specific GCaMP recordings, may help elucidate the dynamics of different classes of dopamine neurons and their implications for behavior.

## STAR Methods

### Key Resources Table

**Table 1.**

| REAGENT or RESOURCE | SOURCE | IDENTIFIER |
| --- | --- | --- |
| **Viral strains** |  |  |
| pAAV-hsyn-GRAB_DA2h | Addgene | 140554-AAV9 |
| pENN.AAV.CB7.CI.mCherry.WPRE.RBG (AAV9) | Addgene | 105544-AAV9 |
| **Experimental models: Organisms/strains** |  |  |
| Long-Evans Rats | Hilltop Lab Animals | Hla®(LE)CVF® |
| Long-Evans Rat | Charles River | 006 |
| **Software and algorithms** |  |  |
| MATLAB | MathWorks | R2023a, R2024a |
| **Chemicals, peptides, and recombinant proteins** |  |  |
| GFP Polyclonal Antibody | Thermo Fisher Scientific | A11122 |
| Goat anti-Rabbit IgG (H+L) Cross-Adsorbed Secondary Antibody, Alexa Fluor™ 488 | Thermo Fisher Scientific | A11008 |
| **Other** |  |  |

### Resource Availability

#### Lead Contact

Further information and requests for resources and reagents should be directed to and will be fulfilled by the lead contact, Christine Constantinople.

#### Materials availability

This study did not generate new unique reagents.

## Data and code availability

### Experimental model and study participant details

A total of 347 Long-evans rats (*Rattus norvegicus*; 215 males, 132 females) between the ages of 6 and 24 months old. We found no effect of age on the main behavioral findings (Supp. Fig. 6). This cohort included 24 TH-Cre rats, 8 ADORA2A-Cre, and 3 DRD1-Cre rats. We also did not find any effect of genotype on the main behavioral findings (Supp. Fig. 7) Animal procedures were approved by the New York University Animal Welfare Committee (UAWC #2021-1120) and carried out in accordance with National Institute of Health standards.

Rats were typically pair-housed. To motivate behavioral performance, rats were water restricted from Monday to Friday, during which time they received water during behavioral training sessions, typically 90 minutes, followed by 20 minutes of ad libitum water. Rats were given ad libitum water following training on Friday though mid-day Sunday. Rats were weighed daily.

### Method Details

#### Behavioral training

We have previously published a detailed description of the behavioral shaping procedure for this task^18^. Rats performed a self-paced temporal wagering task. Rats initiated trials by maintaining a nose poke in the center port for a variable period drawn from a uniform distribution over [0.8, 1.2] seconds. As the rat maintained the nose poke, the reward offer on that trial was conveyed by an auditory tone [1, 2, 4, 8, 16 kHz], which mapped onto one of five rewards ([5, 10, 20, 40, 80*µ*L] for males, [4, 8, 16, 32, 64*µ*L] for females). Following the reward tone presentation, rats could either wait a random delay drawn from an exponential distribution with mean of 2.5 seconds to receive their reward, or could opt-out at any time to immediately start a new trial. On 15-25% of trials (catch trials), reward was withheld to force the rats to exercise the opt-out option.

#### Training for male and female rats

We collected data from both male and female rats (215 males, 134 females). Male and female rats were trained with the same shaping procedure. Early cohorts of female rats experienced the same reward set as the males. However, because female rats are smaller, they consumed less water and performed substantially fewer trials than the males. Therefore, to obtain sufficient behavioral trials from both, females reward offers were slightly reduced while maintaining the logarithmic spacing: [4, 8, 16, 32, 64*µ*L]. For behavioral analysis, reward volumes were treated as equivalent to the corresponding volume for the male rats (e.g., 16*µ*L trials for female rats were treated the same as 20*µ*L trials for male rats). The auditory tones were identical to those used for male rats. We did not observe any significant differences between the male and female rats, in terms of contextual effects, or behavioral dynamics at block transitions^18^. Photometry data in this study was collected from females.

#### Criteria for including behavioral data

To determine whether rats sufficiently understood the mapping between auditory cues and water reward volumes, we evaluated their wait times on catch trials as a function of offered rewards. For each session, we first removed wait times that were greater than two standard deviations from the mean, which likely reflected lapses in attention/task disengagement. Next, we regressed wait time against offered reward. We included sessions with significant positive slopes that preceded at least one other session with a positive slope. We excluded trials with trial initiation times above the 99th percentile of the rat’s cumulative trial initiation time distribution pooled over sessions.

#### Behavioral modeling

To model trial initiation times, we developed computational models based on [18, 68], which describe the optimal trial initiation time, *TI*, given the value of the environment, *V*, as

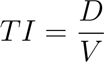

where *D* is a scale parameter. We developed multiple computational models that instantiated different algorithms for estimating the value of the environment. We estimate the value of the environment on trial *t*, *V_t_*, by the recursive formula

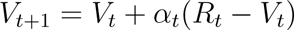

where *α_t_* = *g_t_ · α*_0_ is the learning rate, *g* is the learning rate gain, and *R_t_* is the log_2_(reward) on the current trial. For the static learning rate model, *g* = 1 for all trials.

#### Dynamic learning rate models

We tested several models of dynamic learning rates.

##### 1. Mackintosh surprise model

In this model, the gain on the learning rate is proportional to the salience of that trial^19^, which we assumed to be directly proportional to the reward offer volume on that trials so,

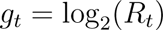

where *α*_0_ is the base learning rate.

##### 2. Pearce-Hall model

In this model, the learning rate gain is directly proportional to the inferred volatility of the environment. Volatility in this model is “model-free” as estimated as the unsigned RPE on the previous trial^22^, so

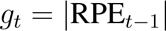

where RPE*_t−_*_1_ is the reward prediction error on the previous trial.

##### 3. Δ Belief model

In this model, as in 2, the learning rate is directly proportional to the inferred volatility of the environment. In this model, volatility is calculated using the trial-by-trial change in the belief of being in a mixed block, using Bayes rule and knowledge of the underlying block structure^18^, so

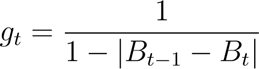

where *B_t_* = *P* (Block = Mixed *| R_t_*). We used the mixed block probability as a summary statistic for the full posterior distribution over blocks, as there is always some ambiguity about whether the animal is in a mixed block, and the block probabilities all need to sum to one. Therefore, changes in the probability of being in a mixed block reflect changes in the full posterior distribution on each trial^18^.

#### Fitting and evaluating models

We fit the models by minimizing the negative-log-likelihood of the the model using MAT-LAB’s constrained minimization function, *fmincon*, assuming log-normal noise with constant variance (variance = 1.7, selected from cross-validated grid search on a subset of rats). We used 100 random seeds and selected the fit with the lowest negative-log-likelihood. We have previously validated our fitting procedure by fitting the models to generative datasets with known parameters^18^. We used 5-fold cross-validation to fit five sets of parameters to each rat (one for each fold), and selected the parameters with the lowest negative log-likelihood per trial on that fold’s test set. Finally, we evaluated the performance of the model fits on a final held-out validation set of trials.

We fit the static learning rate model to the rats’ trial initiation times in early and late trials separately. From previous work, sequential learning effects were primarily driven by post-violation trials^18^, so we fit the model to only post-violation trials. Furthermore, the distribution of trial initiation times was generally heavy-tailed, and seemed to reflect multiple processes on different interacting timescales (e.g., reward sensitivity on short timescales, attention, motivation, and satiety on longer timescales). To capture only task-engaged trials, we removed trial initiation times above the 90th percentile of trial initiation times pooled over sessions for each rat.

#### Bayesian Online Changepoint Detection model

We compared the dynamic learning rate models to a normative Bayesian online changepoint detection model^23^. This model identifies abrupt changes, or changepoints, in the underlying generative distribution of sequentially observed data, which in our case corresponds to block transitions. The time between changepoints is called the run-length. On trial *t*, the model looks at the last *N* trials (*N* ranges from 0 to *t*) and estimates the probability that these *N* observations come from a different distribution than the trials before them. If that probability is high, then the model returns a run-length of *N*, meaning a changepoint occurred *N* trials ago.

Let *x_t_* denote the observation on trial *t* and **x**_*t*_1:_*t*_2__ be the sequence of observations from *t*_1_ to *t*_2_, inclusive, i.e., {*x*_*t*_1__ *, x*_*t*_1_+1_*, …, x*_*t*_2__*_−_*_1_*, x*_*t*_2__ }. On trial *t*, the run-length, *r_t_* can range from 0 to *t*. Finally, given a run-length, *r_t_*, let **x**_*t*_^(*r_t_*)^ be the observations since the last changepoint, that is, **x**_*t−r_t_*:*t*_. We calculate the probability of each potential run-length, known as the run-length posterior, with

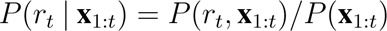

We can simplify the above by marginalizing over the previous run-lengths, *r_t−_*_1_, applying the chain rule, and the assumptions that our data are independently generated, giving us

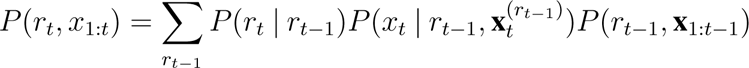

The first term, *P* (*r_t_ | r_t−_*_1_) is called the changepoint prior and captures how often change-points occur, which depends on the hazard rate. Given a previous run-length, *r_t−_*_1_, the next run-length can only be *r_t−_*_1_ + 1 (a changepoint did not occur) or 0 (a changepoint did occur). As described in [18], for simplicity, we assume that the hazard rate is constant with a value of 1*/*40, so we have

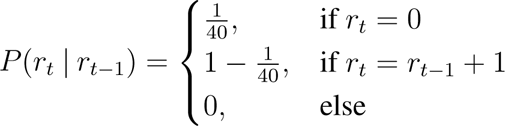

The second term, *P* (*x_t_|r_t−_*_1_, **x**_*t*_^(*r*_*t*−1_)^), is called the predictive probability. This term calculates the high level intuition given above: given some hypothetical run-length, are the data since that run-length consistently from one distribution. To calculate this, we assume that the rats have knowledge of the underlying block structure. We can calculate the predictive probability by marginalizing over the blocks, *B*, giving us

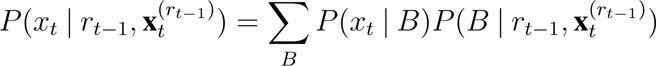

The first term is simply the likelihood of *x_t_* given a block. We can use Bayes rule to calculate the second term, giving us

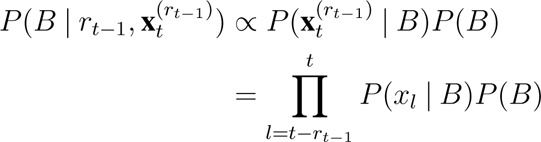

which calculates the likelihood that each datapoint since the hypothetical changepoint belongs to each of the three blocks, and weights by the prior for that block. For simplicity, we assume that the block prior is constant and flat, meaning *P* (*B*) = 1*/*3 for all blocks.

The final term, *P* (*r_t−_*_1_, **x**_1:_*_t−_*_1_) is simply the posterior from the previous trial, so we can recursively update the posterior using the estimate from the previous trial, multiplied by the changepoint prior and the predictive probability, appropriately normalized. The probability of a changepoint was defined as the probability density at *r_t_* = 1, that is, the probability that a changepoint just occurred.

On each trial, the number of computations grows linearly for each trial, so the model has time complexity *O*(*N* ^2^), meaning that doubling the number of trials roughly quadruples the number of computations, which can become costly for long sessions. For this paper, following [23], we implement a modified version that only calculates run-lengths *<* 75 trials. This modification allows us to run the model with constant time complexity, *O*(0), and returns essentially equivalent results as the full model since changepoints occur every 40 trials and thus we do not expect potential run lengths *>* 75. It is worth noting, however, that this truncated implementation still requires 75 computations per trial and requires remembering the previous 75 rewards in order, so is still unlikely to be feasible for rats to be performing.

#### Sterotaxic surgeries

We performed all surgeries using a Neurostar Robot Stererotaxic system on rats after 4 months of age. All rats were induced with 3% isofluorane in oxygen at a flow rate of 2.5 L/minute, which was reduced to 2% isofluorane in oxygen at a flow rat of 1.75 L/minute for maintenance for the duration of the procedure. NAcc injections and implants were targeted to AP 1.3; ML 1.65; DV -6.9 with an 8-10*^◦^* angle from the midline for bilateral implants.

#### Photometry

We measured dopamine release using fiber photometry and GRAB_DA_ sensors (AddGene #140554). We injected AAV9-hsyn-GRAB DA2h to drive expression of the GRAB sensor, as well as AAV1-CB7-CI-mCherry-WPRE-RBG (AddGene #105544) to drive the expression of mCherry to correct for motion artificats. Rats received 60 nL, both delivered over a range of DV values. We implanted 400*µ*m, 0.5 NA chronically implantable optic fibers (Thorlabs) over the injection site (DV -6.7 - -6.9). We simultaneously recorded GRAB_DA_ and mCherry fluorescence with Doric Lenses hardware and software (Doric Neuroscience Studio).

We preprocessed the data and corrected for motion using Two-channel Motion Artifict Correction (TMAC)^69^. First, slow changes in the DC signal due to photobleaching over time were removed by subtracting an exponential decay fit to the session. Next, TMAC removed motion artifacts from the GRAB channel using the control fluorescent channel (either mCherry of isosbestic recordings of GFP). Briefly, TMAC subtracts motion artifacts inferred from the control channel, while accounting for statistically independent sources of noise in both channels. For a subset of rats, we corrected for motion artifacts using both the mCherry signal as well as isosbestic recordings of GFP. We found similar results for both methods. Finally, individual sessions are z-scored using the entire sessions mean and standard deviation (Supp. Fig. 8).

### Quantification and Statistical Analysis

#### Sensitivity to reward blocks

To assess sensitivity to blocks across the population, we z-scored each rat’s trial initiation time using the cumulative mean and standard deviation pooled across sessions, and averaged z-scored trial initiation times over blocks. For the example rat, we compared the median trial initiation time pairwise for each possible pair of blocks using a Wilcoxon sign-rank test. Across the population, we compared average trial initiation time for each pair of blocks using a paired Wilcoxon sign-rank test.

#### Block transition dynamics

To examine how behaviors changed around block transitions, for each rat, we z-scored their trial initiation times. We removed satiety effects by regressing trial initiation times against trial number and subtracted the fit. We then averaged the z-scored trial initiation times based on their distance from a block transition, including violation trials (e.g., averaged all trials five trials before a block transition). Finally, for each transition type, we smoothed the average transition curve using a causal filter (in order to not introduce pre-transition artifacts) of 10 trials individually for each rat. Finally, we averaged transition curves across rats for each transition type.

#### Previous reward regression

To capture the trial history effects, we regressed trial initiation time against previous rewards. We focused on mixed blocks only. We linearized the rewards by taking the binary logarithm of each reward, log_2_(reward), and set the reward for unrewarded trials (e.g., violation or catch trials) to 0, since rats do not receive a reward on those trials. We regressed the previous nine log_2_(reward) offers, not including the current trial, with a constant offset using Matlab’s builtin regress function. We set the first non-significant coefficient (coefficient whose 95% confidence interval overlapped with 0) and all subsequent coefficients to 0. To quantify the time scale of the coefficients, we fit a negative exponential decay curve of the form coefficient*_t_* = *D* exp(*−x/τ*) to each rat’s previous trial coefficients, and reported the time constant (*τ*) for each rat. If rats had one or fewer significant previous trial coefficients, tau was reported as NaN. For early and late block regressions, we used an identical procedure, but only on the first or last 10 trials of a mixed block. To assess the number of significant previous coefficients, for each regression coefficient, we compared the population median coefficient to 0 using a Wilcoxon signed-rank test. To compare *τ* early and late *τ* fit to the regression coefficients, we used a paired Wilcoxon Sign-rank test across the population.

#### Photometry

For all photometry analyses, to quantify dopamine release, we measured the AUC of the dopamine response by integrating the dopamine fluorescence from 0 to 0.5 seconds from the event alignment. DA signals were not baseline corrected with the exception of Figure 4M. In that case, for each trial, baseline was defined as the average response from 0.5 to 0 seconds before delay end, which was subtracted out from that trial. Except where noted, all dopamine analyses were restricted to mixed blocks. To assess reward history effects on NAcc dopamine fluorescence, we used similar methods as above, with the inclusion of an additional coefficient for the current trial offer. To compare dopamine AUC to model estimates of reward prediction error for early and late trials (first or last 10 trials), we used the RPE estimates fit to the respective trial type and the dopamine responses only on those trials. Then for each individual session, we regressed the NAcc dopamine response against RPEs separately for positive and negative RPEs, given the rectification of negative RPE encoding, using Matlab’s builtin robustfit function.

## Acknowledgments

We thank members of the Constantinople lab for feedback and helpful discussions. We thank research technicians in the Constantinople lab for animal training.

Funding: This work was supported by R00MH111926, an Alfred P. Sloan Research Fellow-ship, a Klingenstein-Simons Fellowship in Neuroscience, and an NIH Director’s New Innovator Award (DP2MH126376) to C.M.C. C.G. was supported by a grant from the Simons Foundation (855332)), F32MH125448, and 5T32MH019524. A.M. was supported by 5T90DA043219, F31MH130121, and 5T32MH019524.

## Author Contributions

C.M.C. and A.M designed the study. C.E.G. collected the photometry data and contributed to photometry analysis. A.M. developed the behavioral model and performed behavioral and photometry analyses. A.M. prepared the figures. C.M.C. and A.M. wrote the manuscript. C.M.C. supervised the project.

## Supplemental Material

**Supp. Fig.1:**
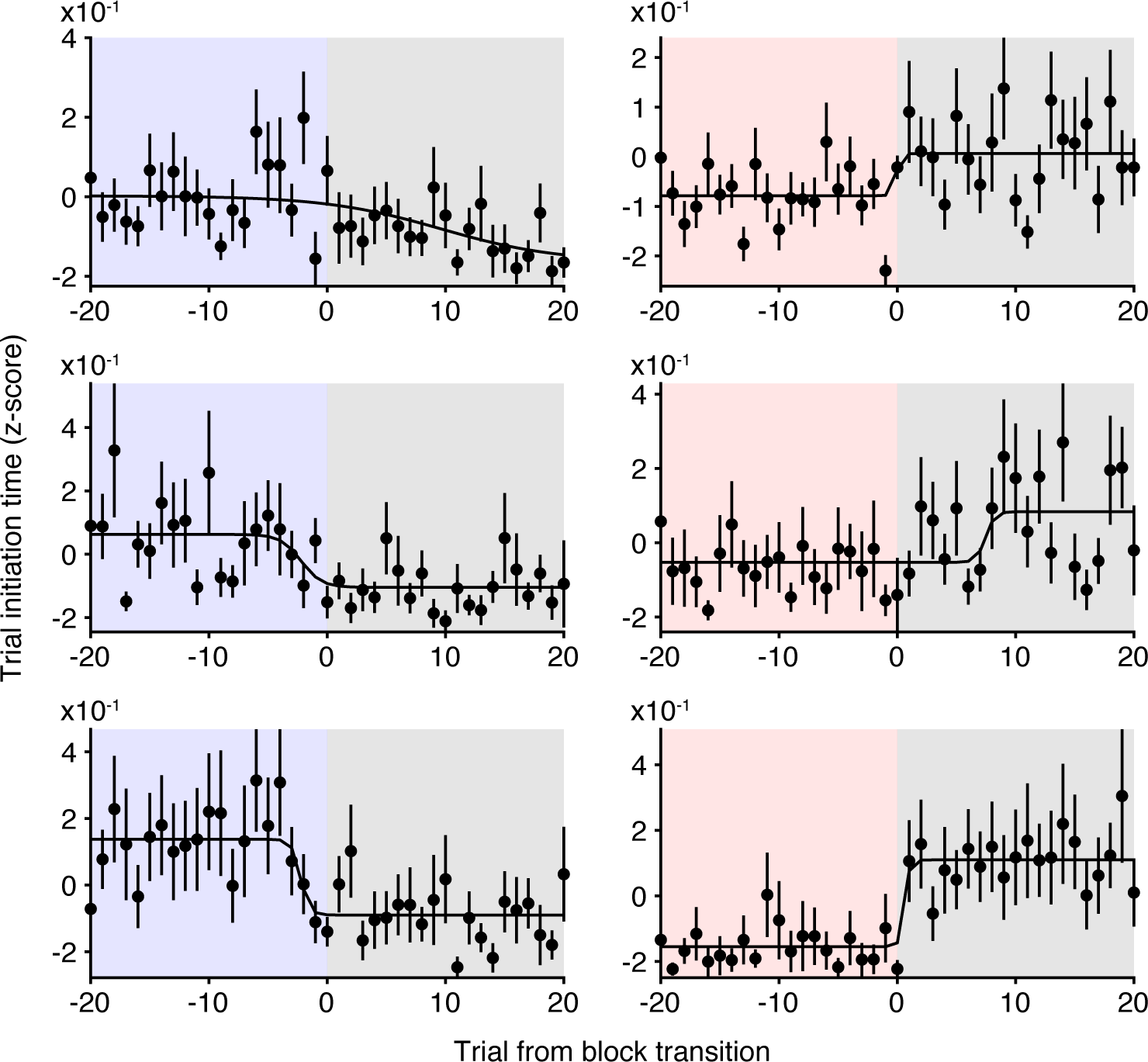
Raw trial initiation time block transition plots for three example rats. Solid lines are sigmoid function fit to the average data. Data are mean *±* S.E.M.

**Supp. Fig.2:**
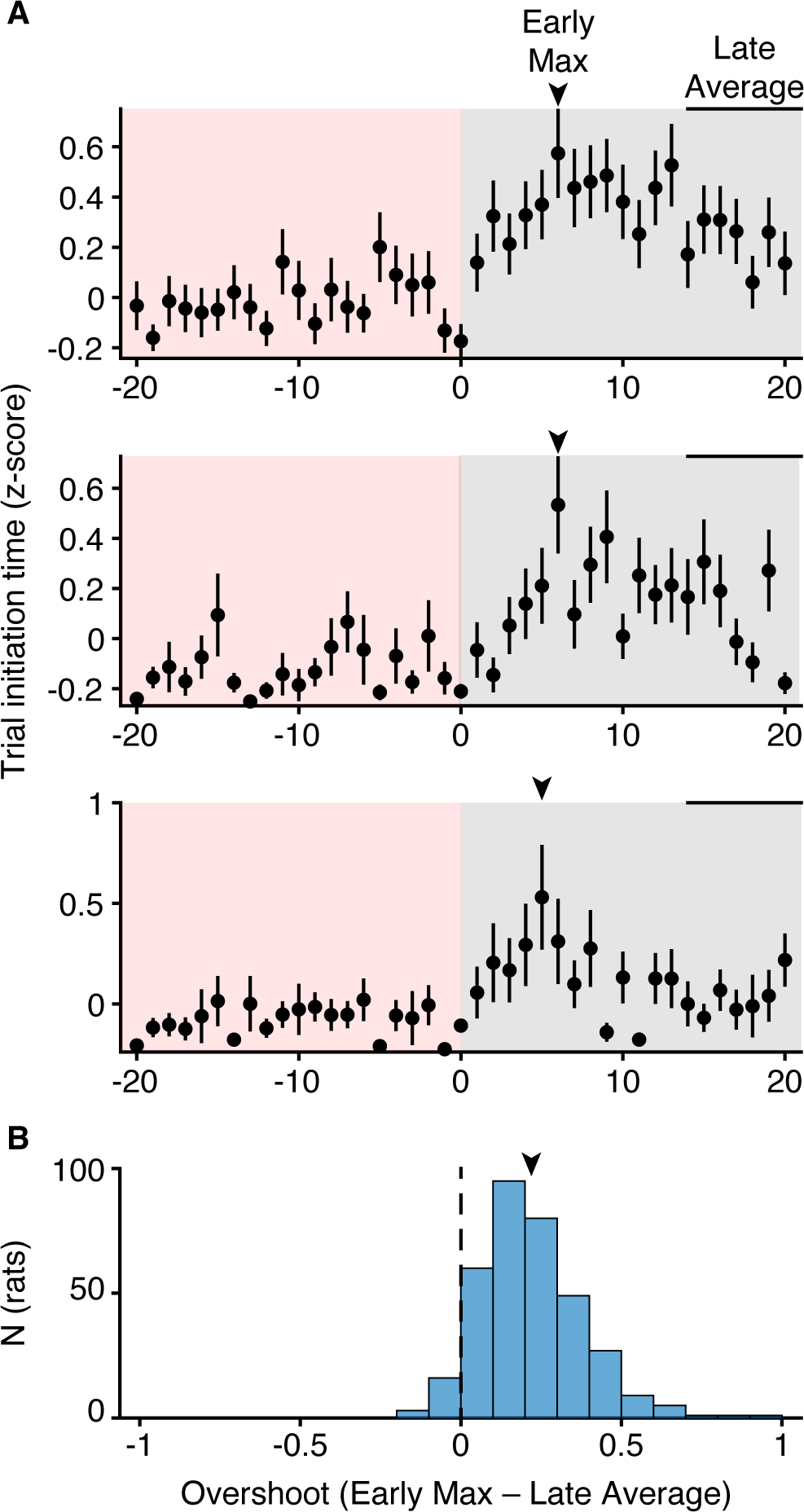
Overshoot is robust across rats. **A**. Three example rats showing trial initiation time overshoot. Arrow indicates maximum early trial initiation time and bar indicates window for average to calculate overshoot in B. **B.** Overshoot quantified across rats. Arrow indicates mean. All data are mean*±*S.E.M.

**Supp. Fig.3:**
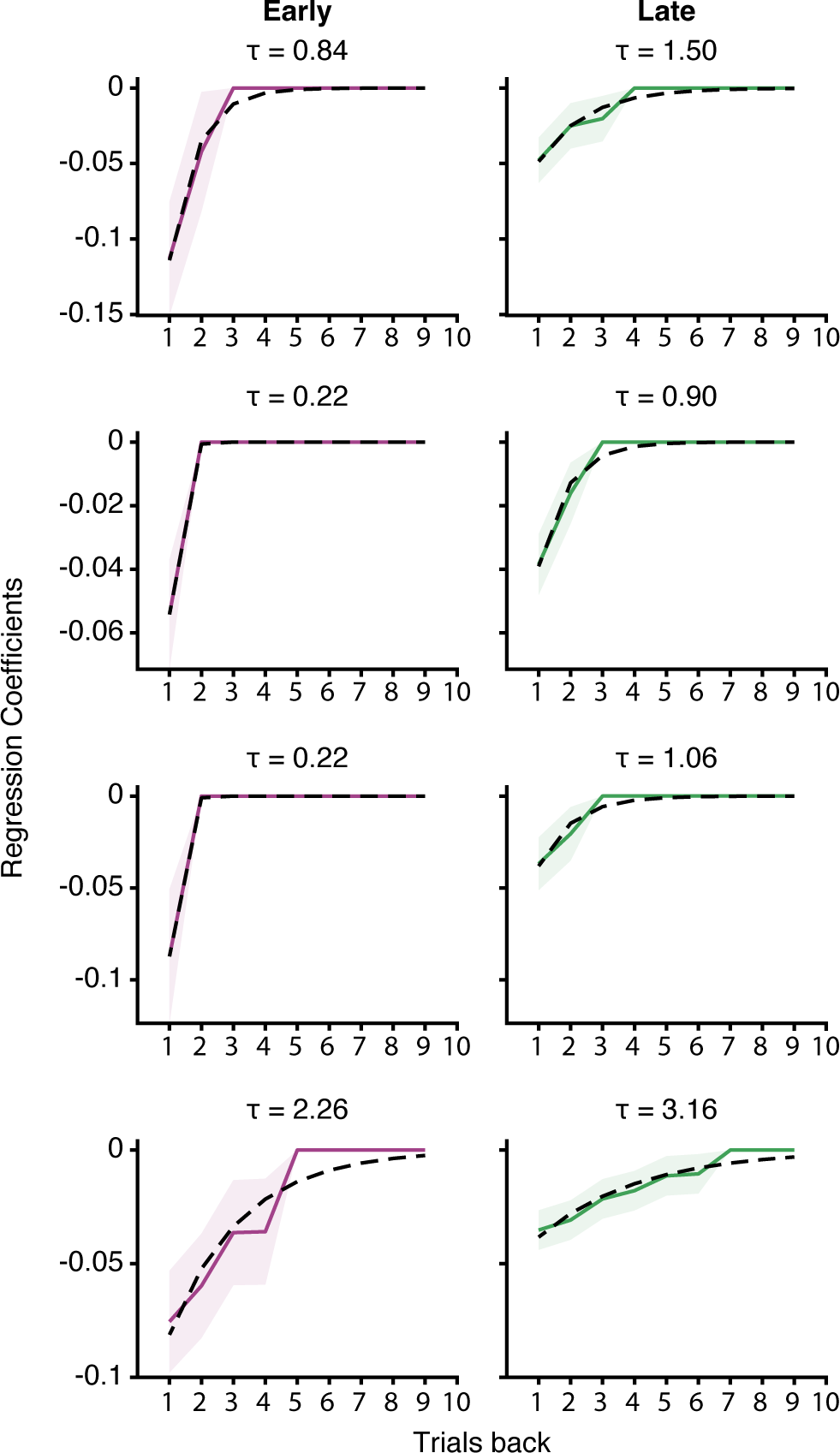
Previous reward regression coefficients for (left) early and (right) late mixed block trials for 4 example rats. Solid line are coefficients, dashed are exponential curve fit to coefficients. Title indicates time constant of fit exponential. Data are mean *±* 95% confidence interval

**Supp. Fig.4:**
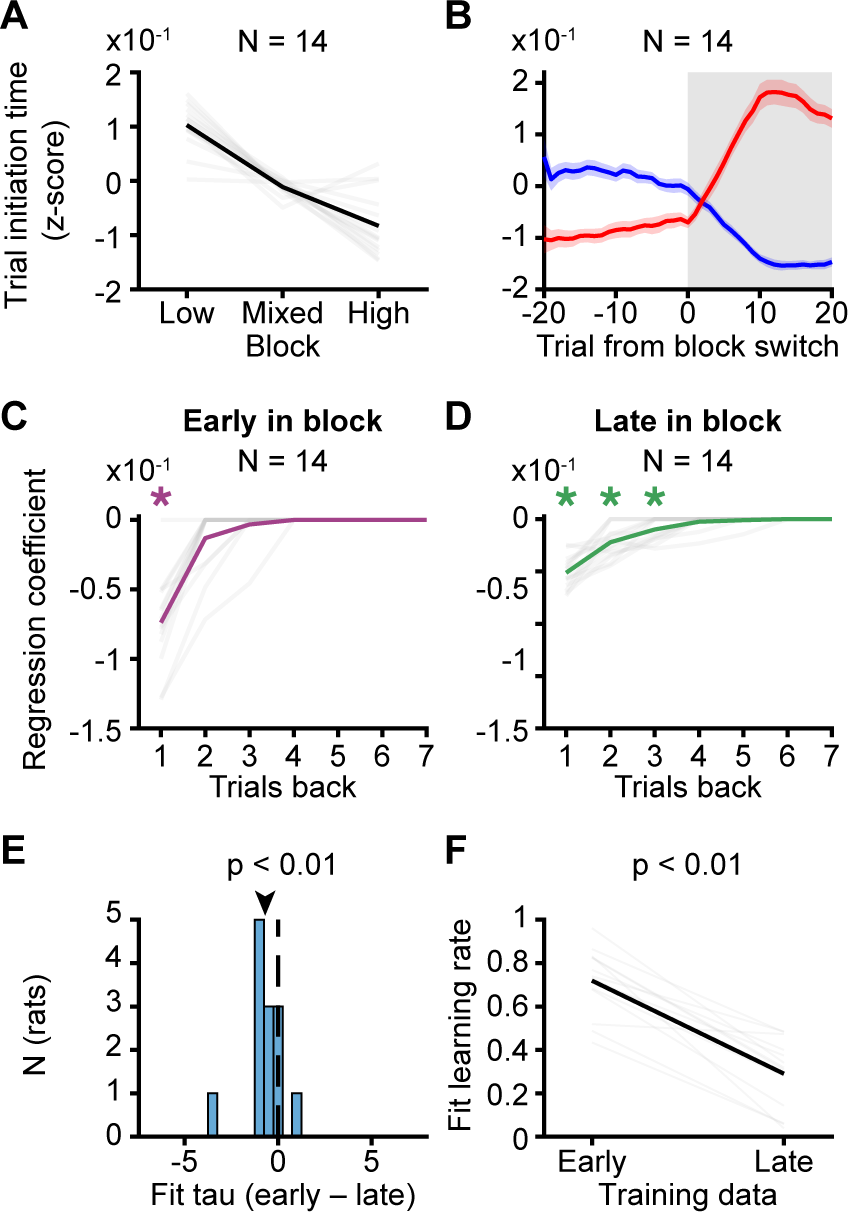
Fiber photometry rats have similar behavior to full population. **A.** Average trial initiation times by block. **B.** Trial initiation times aligned to transitions into mixed blocks from low (blue) and high (red) blocks. Smoothed with a causal filter of size 10 trials. **C-D.** Previous reward regression coefficients for (C) early and (D) late mixed block trials (Wilcoxon sign-rank test, *N*=14). **E.** Difference in time-constant, tau, of exponential decay fit to early and late mixed block regression coefficients across rats. Arrow indicates mean (*p<* 0.01, paired Wilcoxon Sign-rank test, *N*=14). **F.** Recovered learning rate parameters for early and late mixed block training data across rats (*p<* 0.001, Wilcoxon Sign-rank test, *N*=14).

**Supp. Fig.5:**
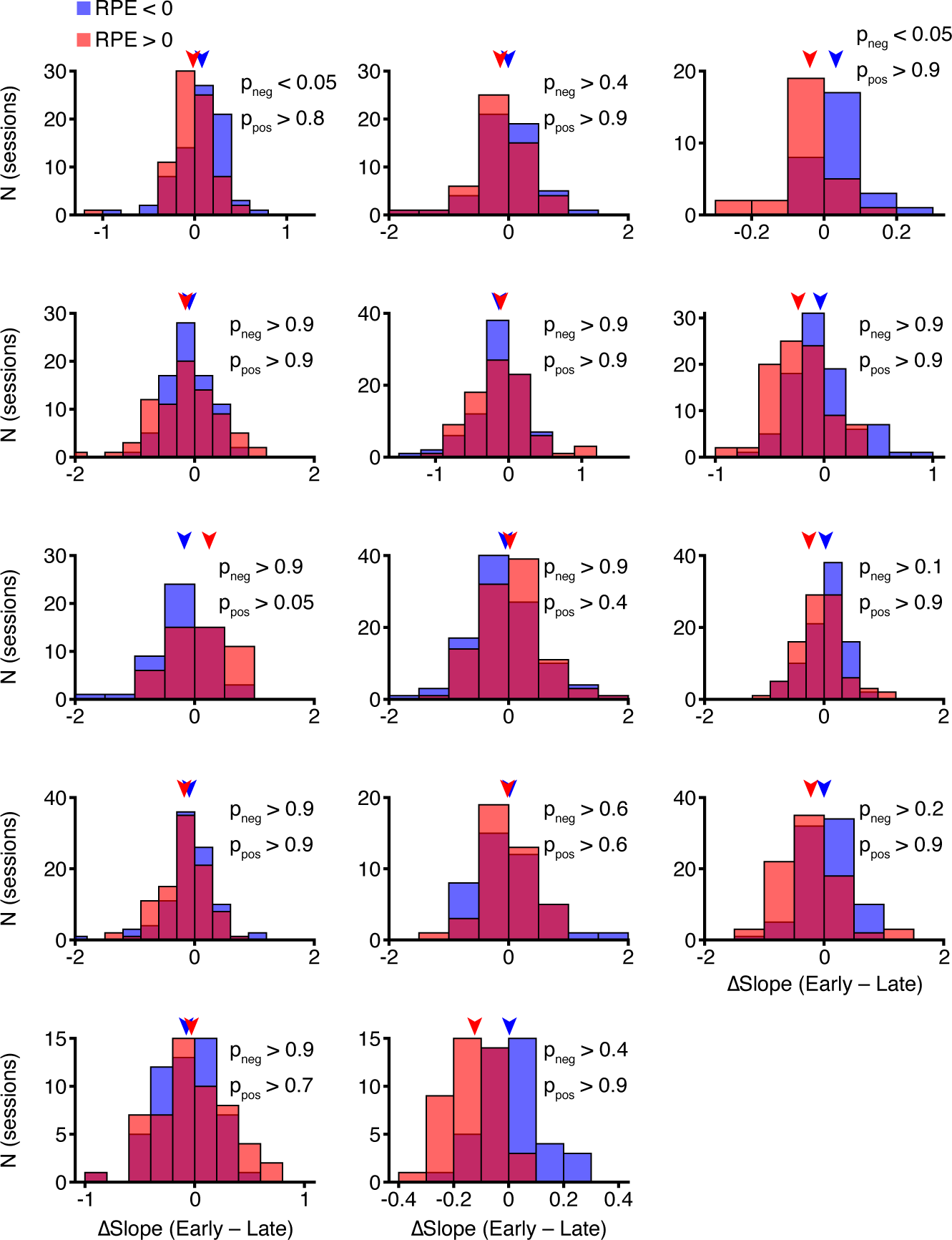
RPE vs. DA slopes are comparable early and late in mixed blocks across photometry rats. Change in slope for early vs. late DA vs. RPE regression, fit separately for positive and negative RPEs for each sessions, for individual rats. Two rats had significantly higher slopes for negative RPEs, which is reduced to one after correcting for multiple comparisons. Arrows indicate mean.

**Supp. Fig.6:**
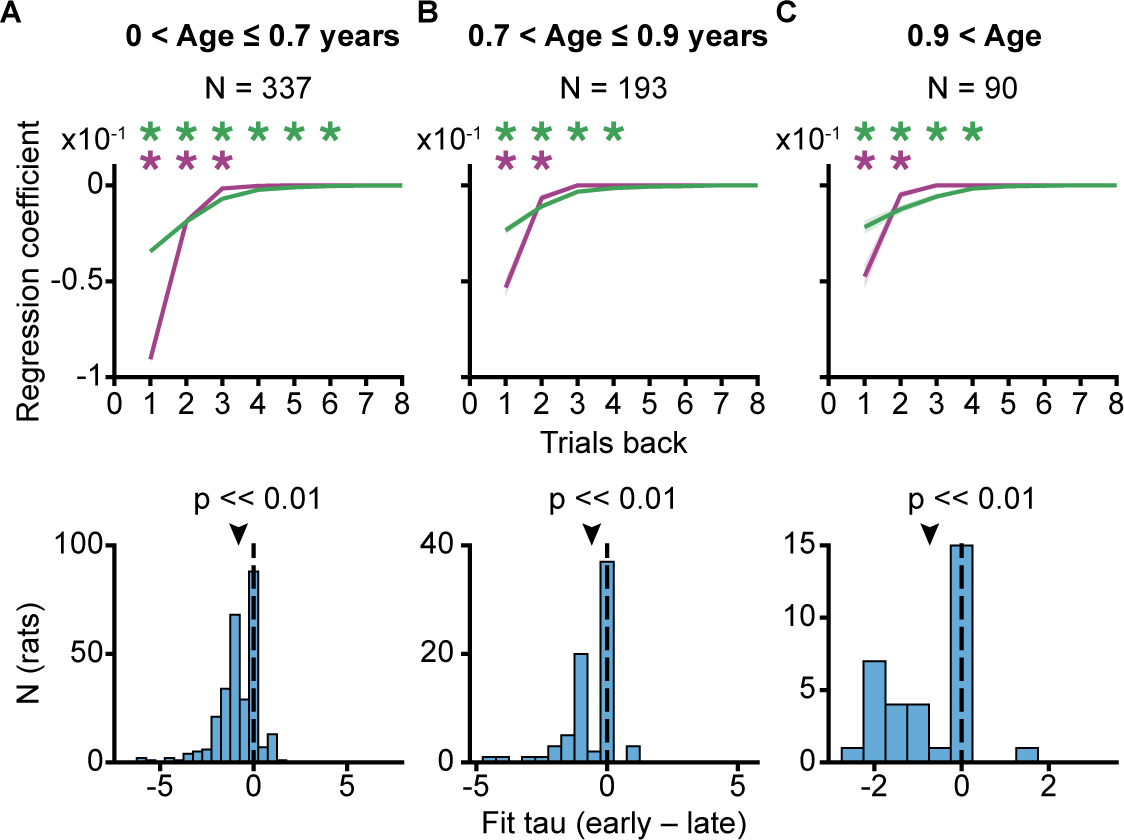
Regression analysis is qualitatively similar across age groups. (Top) Regression analysis separated by age of rat during that session. (A) Age less then 0.7 years old (*N*=337), (B) Age between 0.7 and 0.9 years old (*N*=193) and (C) Age older than 0.9 years old. (Bottom) Difference in time-constant, tau, of exponential decay fit to early and late mixed block regression coefficients across rats. Arrow indicates mean (*p<* 0.01, paired Wilcoxon Sign-rank test). Data are mean *±* S.E.M. Asterisk indicates *p <* 0.05, Wilcoxon sign-rank test.

**Supp. Fig.7:**
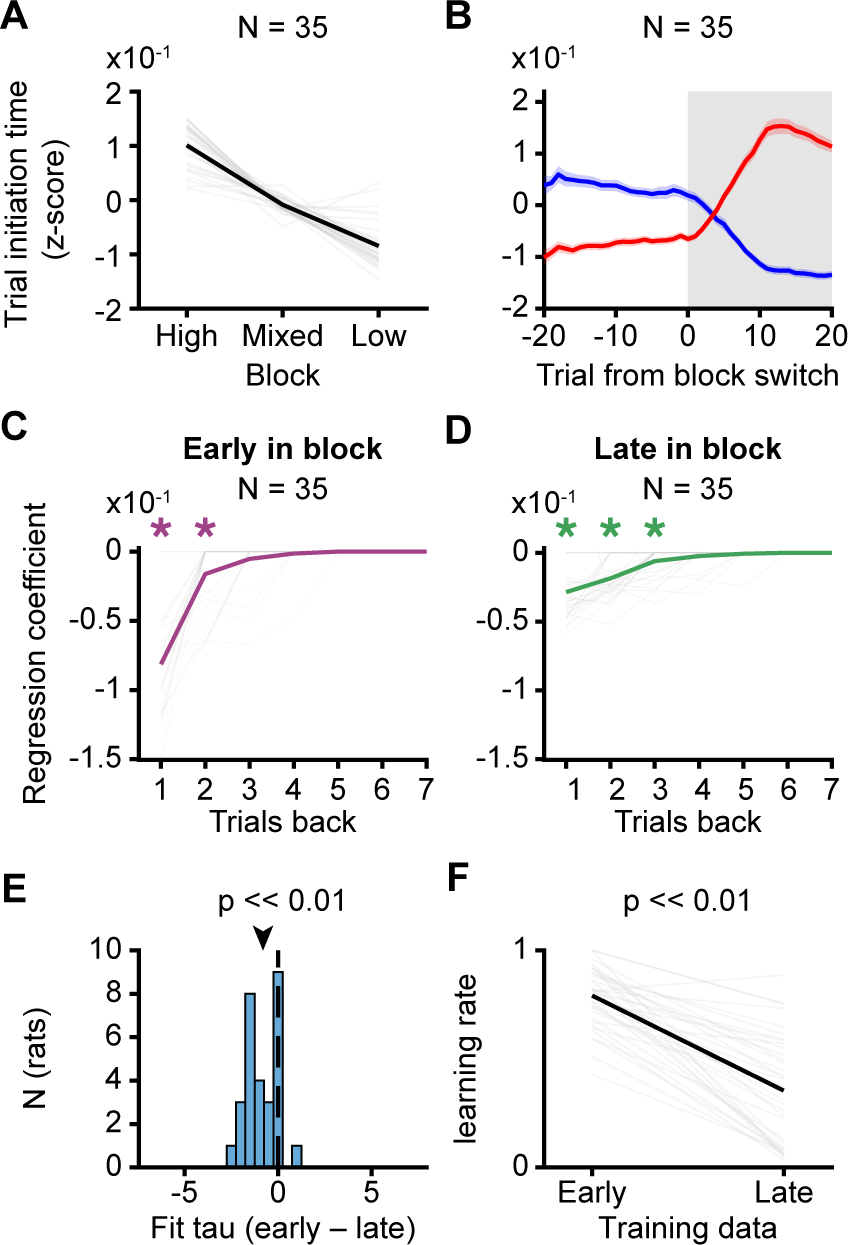
Cre-line rats have similar behavior to full population. Combining behavior across Cre-line rats (24 TH-Cre rats, 8 ADORA2A-Cre, and 3 DRD1-Cre rats) **A.** Average trial initiation times by block. **B.** Trial initiation times aligned to transitions into mixed blocks from low (blue) and high (red) blocks. Smoothed with a causal filter of size 10 trials. **C-D.** Previous reward regression coefficients for (C) early and (D) late mixed block trials (Wilcoxon sign-rank test, *N*=35). **E.** Difference in time-constant, tau, of exponential decay fit to early and late mixed block regression coefficients across rats. Arrow indicates mean (*p<* 0.01, paired Wilcoxon Sign-rank test, *N*= 35). **F.** Recovered learning rate parameters for early and late mixed block training data across rats (*p<* 0.001, Wilcoxon Sign-rank test, *N*=35).

**Supp. Fig.8:**
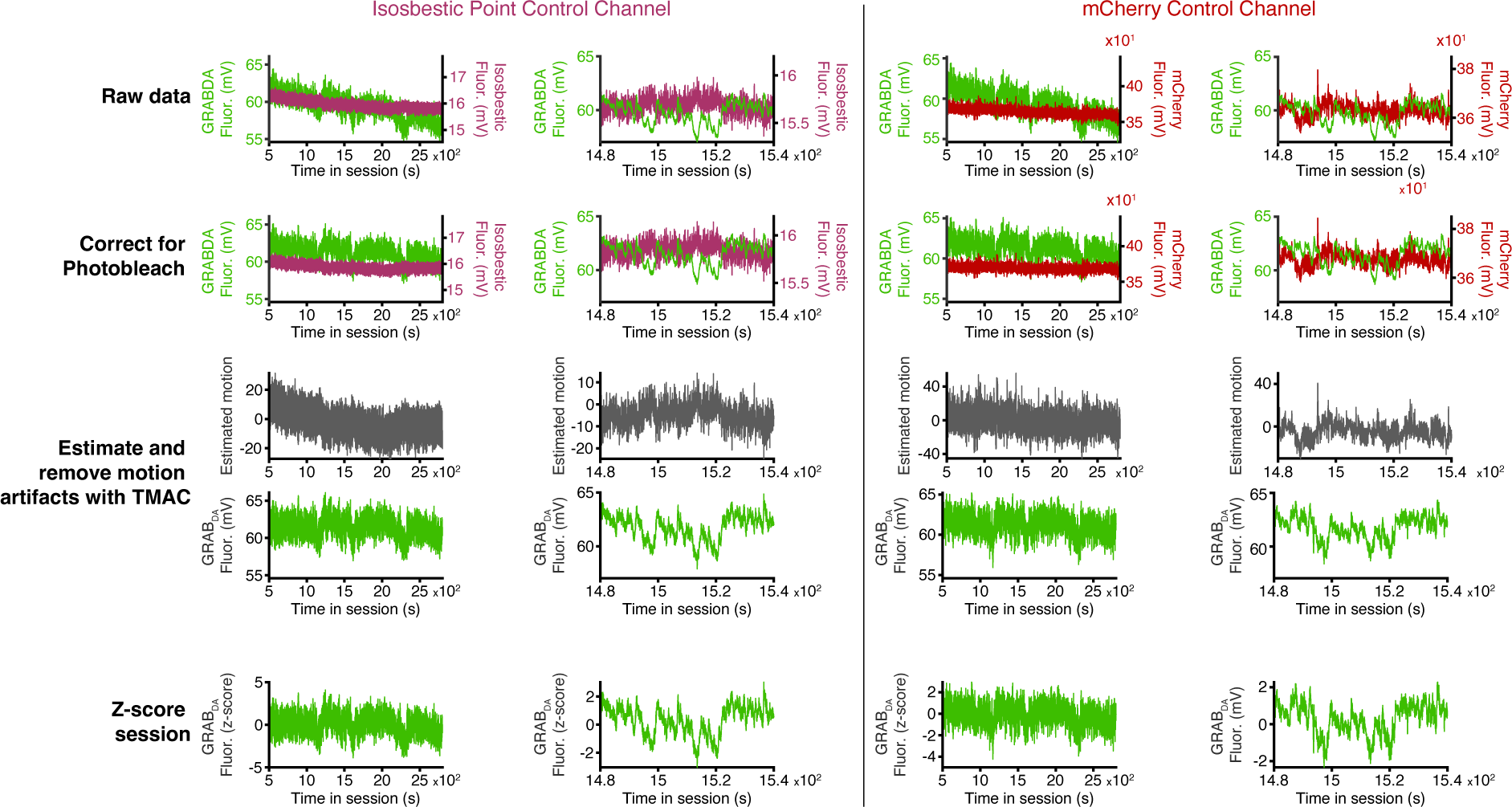
Photometry processing pipeline on an example session with both isosbestic point control and mCherry control channels. For each channel, left column is the full session is a full session, and right is a zoomed in time window. Labels of left indicate step of procedure.

